# Digital pathological classification of peripheral neuroblastic tumors across multi-continent cohorts

**DOI:** 10.64898/2026.10.26.761046

**Authors:** Yuanning Zheng, Yuichi Shibui, Balint Laszlo Forgo, Arlene Naranjo, Lisa Werr, Janina Fischer-Mertens, Matthias Fischer, Naonori Kawakubo, Yoshinao Oda, Tatsuro Tajiri, Mark A. Applebaum, Hiroyuki Shimada, Olivier Gevaert, Bill Chiu

## Abstract

Peripheral neuroblastic tumors (pNTs) are embryonal tumors of neural crest origin with highly variable clinical behaviors, ranging from spontaneous regression to aggressive and fatal disease. The International Neuroblastoma Pathology Classification (INPC) defines four diagnostic categories and stratifies tumors into Favorable Histology (FH) or Unfavorable Histology (UH) group based on age-linked assessment of tumor differentiation/maturation and the Mitosis-Karyorrhexis Index (MKI). Prognostic distinction according to INPC offers significant prediction of clinical outcomes, with patients harboring FH tumors demonstrating longer overall and event-free survival than those with UH tumor. Here, we assembled the largest multi-continental whole-slide histology imaging dataset of pNTs to date, comprising 2,030 tumors from the United States, Canada, Australia, New Zealand, Germany and Japan. Using this dataset, we developed NB-PathAssist, a computational pathology framework for automated INPC-based classification from digitized whole-slide images. NB-PathAssist integrates foundation-model-based feature extraction with an archetype-guided SummaryMixing architecture that compresses whole-slide morphology into representative histologic tokens for tumor-level classification. To automate MKI assessment, we manually annotated 14,049 nuclei, classifying them as mitotic-karyorrhectic (MK) tumor cells, non-MK tumor cells, or non-tumor cells. The resulting model was applied to phenotype cells in whole-slide images across 1,649 tissue sections. Across geographically and institutionally diverse cohorts, NB-PathAssist reproducibly assessed key diagnostic features and generated prognostic classifications concordant with reference INPC annotations. The average balanced accuracy for classifying tumor categories, grade of neuroblastic differentiation, MKI and prognostic groups achieved 0.84, 0.86, 0.89 and 0.96, respectively. We packaged the NB-PathAssist algorithms into an end-to-end online platform for scalable and standardized AI-assisted assessment of pNTs.

## Introduction

Peripheral neuroblastic tumors (pNTs) of neural crest origin were once described enigmatic due to a wide range of their clinical behaviors including spontaneous regression, differentiation/maturation, and aggressive progression even refractory to intensive therapy. In 1999, the International Neuroblastoma Pathology Classification (INPC) linking histopathologic features to patient prognosis of this disease was introduced (**Table 1**)^1–4^. The INPC distinguishes four categories: 1) Neuroblastoma (Schwannian stroma-poor) - NB, 2) Ganglioneuroblastoma, intermixed (Schwannian stroma-rich) - GNB-i, 3) Ganglioneuroma (Schwannian stroma-dominant) - GN, and 4) Ganglioneuroblastoma, nodular (composite, Schwannian stroma-rich/stroma-dominant and stroma- poor) - GNB-n. Tumors in the NB category are classified into Favorable Histology (FH) or Unfavorable Histology (UH) group based on the age-linked evaluation of grade of neuroblastic differentiation and Mitosis-Karyorrhexis Index (MKI). Tumors in the GNB-n category are composed of distinct histologic components, including an NB nodule and a GNB-i or GN component. Their prognostic classification as FH or UH group is also determined by the same age-dependent evaluation of grade of neuroblastic differentiation and MKI of the NB nodule^4^. In contrast, GNB-i and GN are always classified as FH and are associated with excellent outcomes, with 5-year event-free and overall survival rates exceeding 95%, reflecting their advanced degree of differentiation/maturation^5,6^. The impact of prognostic distinction by INPC has been continuously tested for over a quarter of century, and its significance was recognized by International Collaboration on Cancer Reporting (ICCR) that adopted INPC to standardize handling and reporting of pNT cases by pathologists worldwide^7,8^.

**Table 1.** International Neuroblastoma Pathology Classification.

| Histology | Subtype | MKI | Age at diagnosis | Prognostic group |
|---|---|---|---|---|
| NB | Undifferentiated | Any | Any | UH |
|  | Poorly differentiated | Low or intermediate | < 18 months | FH |
|  |  |  | ≥ 18 months | UH |
|  |  | High | Any | UH |
|  | Differentiating | Low | < 5 years | FH |
|  |  |  | ≥ 5 years | UH |
|  |  | Intermediate | < 18 months | FH |
|  |  |  | ≥ 18 months | UH |
|  |  | High | Any | UH |
| GNB-i | N/A | N/A | Any | FH |
| GN | N/A | N/A | Any | FH |
| GNB-n | Prognosis group depends on the characteristics of the neuroblastoma nodule |  |  |  |

NB, neuroblastoma; GNB-i, ganglioneuroblastoma, intermixed; GN, ganglioneuroma; GNB-n, ganglioneuroblastoma, nodular; MKI, mitosis-karyorrhexis index; N/A, not applicable; UH: unfavorable histology; FH: favorable histology.

Over the past decade, artificial intelligence (AI) models have demonstrated success in predicting molecular profiles, diagnosis and prognosis from digitized whole-slide histopathology images^9–12^. While the majority of these models have been developed for common adult malignancies, their performance on pediatric cancers remains relatively underexplored. In the context of pNTs, Gheisari et al. combined deep-learning feature extraction protocols with supporter vector machines to classify pNTs from 125 patients into five diagnostic categories^13^. In a more recent study, Ramesh et al. developed attention-based deep learning models to predict the diagnostic category, MKI, and *MYCN* amplification status of pNTs using 172 images, and the models were validated on an independent cohort of 25 images^14^. While these studies provide preliminary evidence of the effectiveness of deep-learning models in the pathology diagnosis of pNTs, the predictive performance is limited by the small size of training datasets. This challenge is further exacerbated by the class imbalance across diagnostic categories: fewer than 10% of tumors are classified as differentiating subtype and only ∼20% exhibit high MKI in the NB category^15,16^. As a result, models trained on small datasets may be biased toward more prevalent tumor subtypes, limiting their robustness and ability to accurately identify less common subtypes.

In this study, we assembled the largest-to-date multi-continent pNT cohort. We combined methodological innovation with cross-platform validation to develop NB-PathAssist, a deep-learning framework for automated diagnosis and prognostic stratification of pNTs from routine hematoxylin and eosin (H&E)-stained whole-slide histology images. In collaboration with pathologists and clinicians, we designed a pathologist-in-the-loop workflow to ensure that model development, predictive targets, and evaluation strategies were aligned with real-world diagnostic practice and established clinical standards. To support MKI assessment, we further generated cell-level annotations for 14,049 nuclei across independent cohorts and trained a dedicated model to identify mitotic and karyorrhectic tumor cells. NB-PathAssist demonstrated robust performance in predicting key histopathologic features, including tumor category, grade of neuroblastic differentiation, MKI, and further prognostic classification distinguishing FH and UH group. To enable the community to use our models, we packaged the NB-PathAssist algorithms into an end-to-end, publicly accessible online platform (https://nb-ai.stanford.edu), where users can upload histology images of pNTs and receive predicted diagnostic categories accompanied by expert-authored text descriptions.

## Results

### Cohort characteristics

A critical requirement for reliable AI models is generalizability across diverse datasets. However, technical variability and batch effects can limit the performance of models trained on data from a single institution or platform, leading to reduced accuracy when applied to external cohorts. This challenge is especially pronounced in rare diseases, where data scarcity further exacerbates variability and hinders robust model development. To ensure the generalizable capacity of our proposed methods, we assembled a multi-continent whole-slide image (WSI) cohort of pNTs, including H&E-stained histology images of 2,030 tissue sections (**Fig.1** and **Table 2**). The dataset was partitioned into (1) a study cohort used for model development, hyperparameter tuning and internal cross-fold validation (n = 1,692) and (2) independent confirmatory cohorts from multi-national origins reserved exclusively for external testing (n = 338).

**Figure 1.**
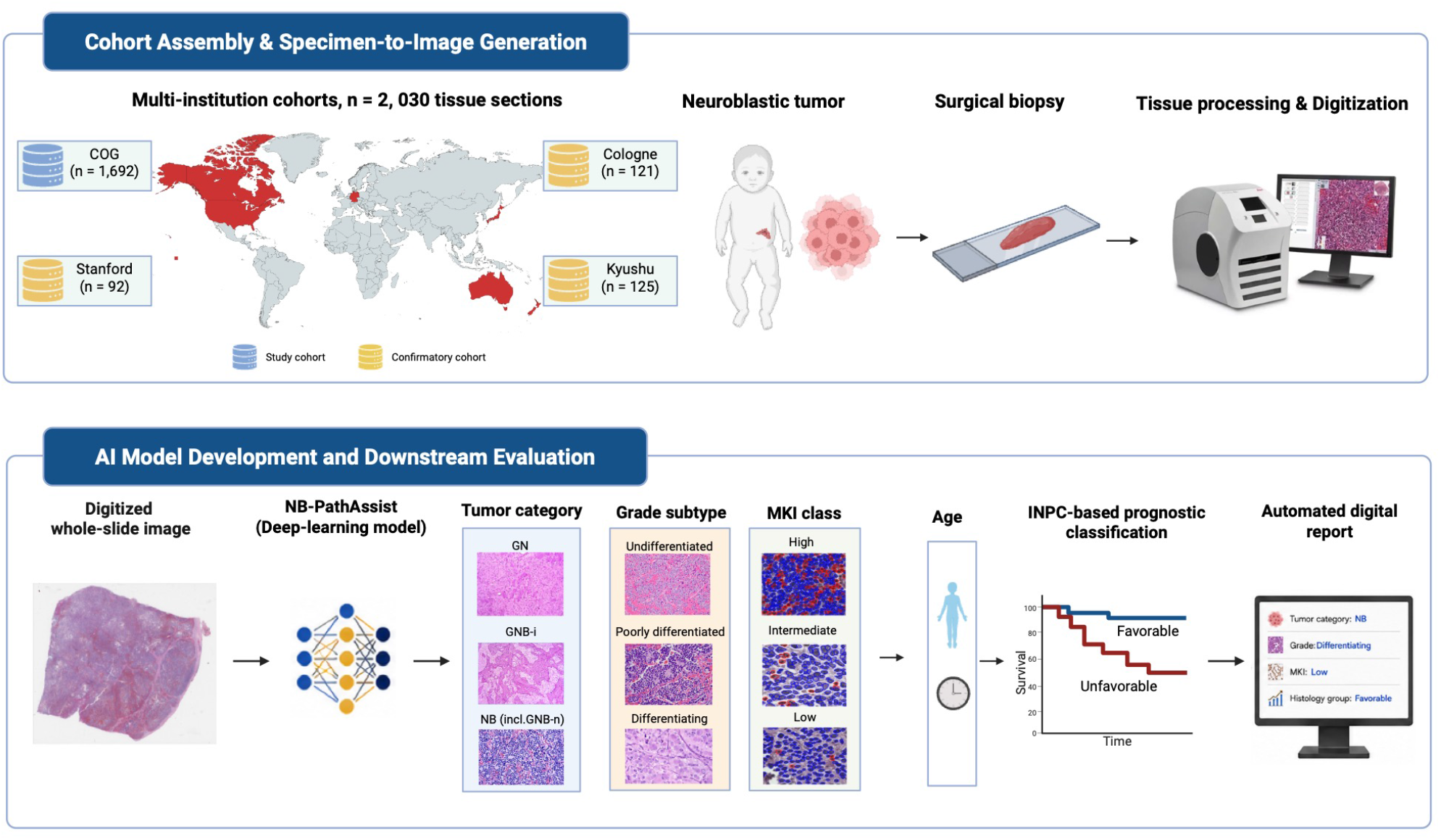
Overview of the study design and patient cohorts. Image was created using BioRender.

**Table 2.** Distribution of peripheral neuroblastic tumor specimens by histological category.

| Cohort | NB | GNB-i | GN | GNB-n | Total |
|---|---|---|---|---|---|
| COG-pNT | 1,264 | 242 | 103 | 83 | 1,692 |
| Stanford-NB | 92 | 0 | 0 | 0 | 92 |
| Cologne-pNT | 102 | 7 | 6 | 6 | 121 |
| Kyushu-pNT | 109 | 7 | 4 | 5 | 125 |
| **Total** | **1,567** | **256** | **113** | **94** | **2,030** |

NB, neuroblastoma; GNB-i, ganglioneuroblastoma, intermixed; GN, ganglioneuroma; GNB-n, ganglioneuroblastoma, nodular; pNT, peripheral neuroblastic tumor.

The study cohort included cases originally collected for the international clinical trials of the Children’s Oncology Group (COG) Neuroblastoma Study with contributions from institutions in the United States, Canada, Australia, and New Zealand. H&E-stained tumor sections were assembled from the COG Neuroblastoma Pathology Reference Laboratory and curated by the creator of the INPC and also director of the Reference Laboratory (H.S.). According to the INPC system, the study cases included tumors in the Neuroblastoma (NB) category (n = 1,264), the Ganglioneuroblastoma, intermixed (GNB-i) category (n = 242), the Ganglioneuroma (GN) category (n = 103), and the Ganglioneuroblastoma, nodular (GNB-n) category (n = 83) (**Table 2**). One representative section from each case was selected by H.S. and Y.S. for whole-slide scanning in this study. For those in the GNB-n category, the selected sections always contained NB nodules whose histologic features were critical for prognostic classification. In this study, the original INPC assessments recorded by H.S. and archived in the COG database, including histologic features and prognostic classifications, were compared with the corresponding predictions generated by NB-PathAssist.

After developing NB-PathAssist, its utility was tested on the confirmatory cohorts including a total of 338 cases from multi-national origins: Stanford Medicine Children’s Health, USA (Stanford-NB, n = 92), Cologne University Children’s Hospital, Germany (Cologne-pNT, n = 121), and Kyushu University Hospital, Japan (Kyushu-pNT, n = 125). All cases of the confirmatory cohorts were also reviewed by HS for this study and the predictive results generated by NB-PathAssist were compared and analyzed.

### Predictive Performance for Tumor Category

We first developed a deep-learning model to distinguish tumors in the NB and GNB-n categories from those in the GNB-i and GN categories using data from the study cohort (**Fig. 2a**). NB and GNB-n were grouped together, designated as NB (incl. GNB-n), because tumors in these categories require further classification as Favorable Histology or Unfavorable Histology based on age-linked assessment of grade of neuroblastic differentiation and MKI (**Table 1**). In contrast, GNB-i and GN are uniformly classified as FH and therefore do not require these additional histologic assessments. To capture representative morphologic patterns across whole-slide images, we developed an archetype-guided SummaryMixing architecture (Archetype-SM; see Methods). Model performance was assessed using ten-fold cross-validation. In each fold, data from 90% of patients were used for model training and hyperparameter tuning, while the remaining 10% were held out for evaluation.

**Figure 2.**
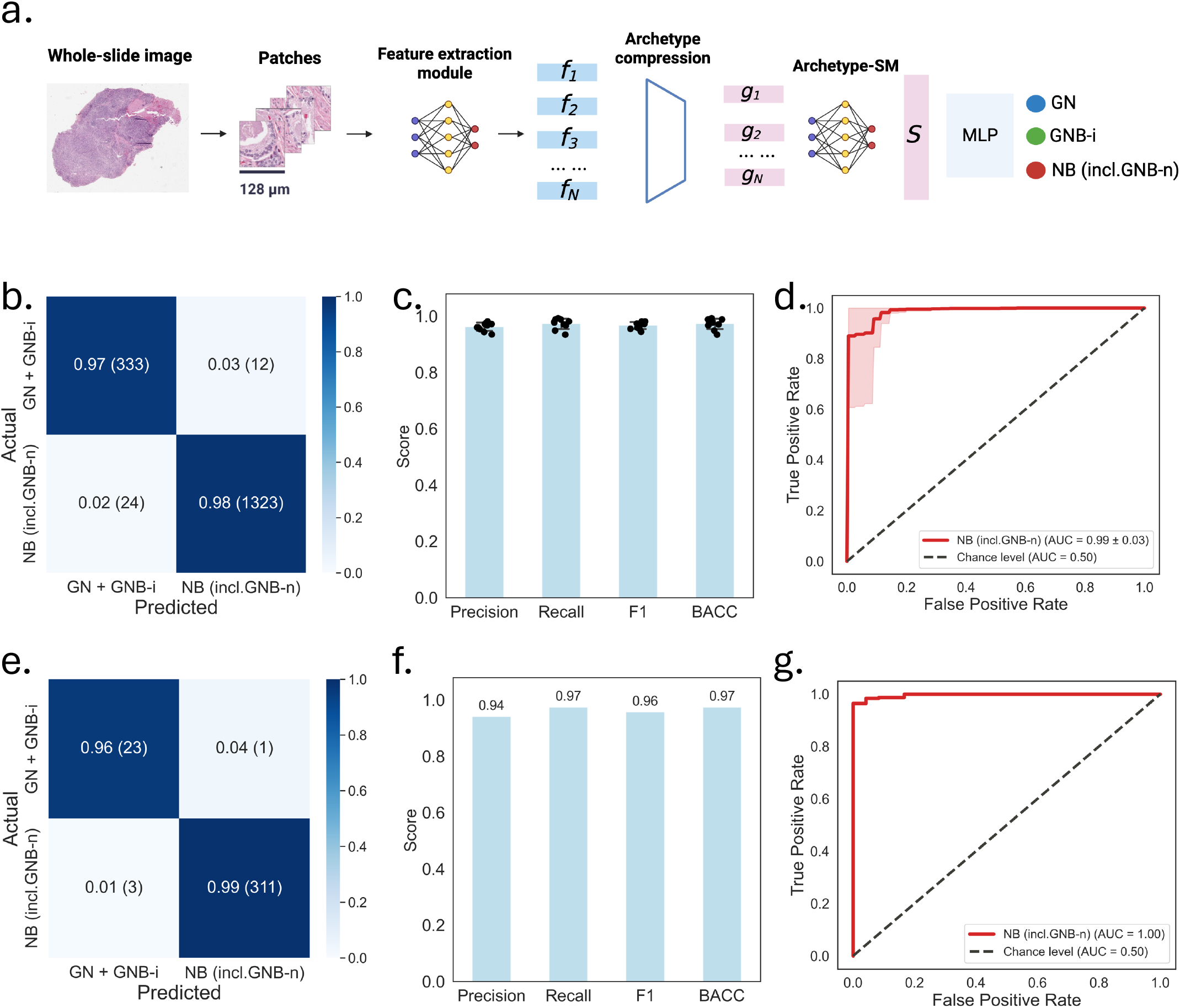
Classification performance for tumor category. **(a)** Overview of the deep-learning architecture. **(b-d)** Classification performance in the COG cohort: (b) confusion matrix; (c) macro-averaged precision, recall, F1 score, and balanced accuracy, with each dot representing the performance of an individual fold. Bars indicate the mean values across the folds, and error bars represent the standard deviation; and (d) receiver operating characteristic curve, with shaded areas representing the standard deviation across cross-validation folds. **(e-g)** Classification performance in the confirmatory cohorts: (e) confusion matrix; (f) macro-averaged precision, recall, F1 score, and balanced accuracy; and (g) receiver operating characteristic curve. NB, neuroblastoma; GNB-i, ganglioneuroblastoma, intermixed; GN, ganglioneuroma; GNB-n, ganglioneuroblastoma, nodular; BACC, balanced accuracy; AUC, area under the receiver operating characteristic curve.

For performance evaluation, we first combined the GNB-i and GN cases into one category and tested whether the model can separate them from the NB (incl. GNB-n) group. Under this binary classification setting, the model achieved a mean macro-averaged precision of 0.96 (standard deviation [SD] = 0.01), recall of 0.97 (SD = 0.02), F1 score of 0.97 (SD = 0.01), and balanced accuracy of 0.97 (SD = 0.02) across all folds (**Figs. 2b-c**). The average area under the receiver operating characteristic curve (AUROC) for identifying the NB (incl. GNB-n) cases was 0.99 (SD = 0.03) (**Fig. 2d**). When GNB-i and GN were considered as separate categories, the model achieved a mean precision of 0.87 (SD = 0.04), recall of 0.89 (SD = 0.04), F1 score of 0.88 (SD = 0.05), and balanced accuracy of 0.89 (SD = 0.04) (**Supplementary Figs. 1a-b**). The per-class AUROC was 0.98 (SD = 0.01) for GNB-i and 0.98 (SD = 0.03) for GN (**Supplementary Fig. 1c**).

We next evaluated model performance using data from the independent confirmatory cohorts from Stanford, Cologne, and Kyushu. Since GNB-i and GN cases represented only a small proportion of all cases (7.1%), we pooled the prediction results across the three institutions for subsequent analysis. The macro-averaged precision, recall, F1 score and balanced accuracy was 0.94, 0.97, 0.96 and 0.97, respectively (**Figs. 2e-f**), and the AUROC was 1.00 (**Figs. 2g**). When GNB-i and GN were considered as separate categories, the model achieved a precision of 0.81, recall of 0.78, F1 score of 0.77, and balanced accuracy of 0.78 (**Supplementary Figs. 1d-e**). The AUROC was 0.98 for GNB-i, and 0.99 for GN (**Supplementary Fig. 1f**). Together, these results demonstrated that our model accurately distinguished the group of NB (incl. GNB-n) from the benign tumor categories GNB-i and GN.

### Predictive Performance for Grade of Neuroblastic Differentiation in the NB and GNB-n Categories

Since prognostic classification of NB and GNB-n tumors depends on age-linked assessment of grade of neuroblastic differentiation and MKI, we next trained a separate model to predict the grade of neuroblastic differentiation using tumors in the NB and GNB-n categories from the study cohort (**Table 3**). The model was designed to classify tumors into undifferentiated, poorly differentiated, or differentiating subtypes. In ten-fold cross-validation, the model achieved a mean macro-averaged precision of 0.70 (SD = 0.05), recall of 0.89 (SD = 0.05), F1 score of 0.74 (SD = 0.06), and a balanced accuracy of 0.89 (SD = 0.05) (**Figs. 3a-b**). Precision was lower for undifferentiated tumors, primarily due to the misclassification of poorly differentiated tumors as undifferentiated (**Fig. 3a**). The mean AUROC was 0.96 (SD = 0.03) for undifferentiated, 0.94 (SD = 0.05) for poorly differentiated, and 0.99 (SD = 0.00) for differentiating tumors (**Fig. 3c**).

**Figure 3.**
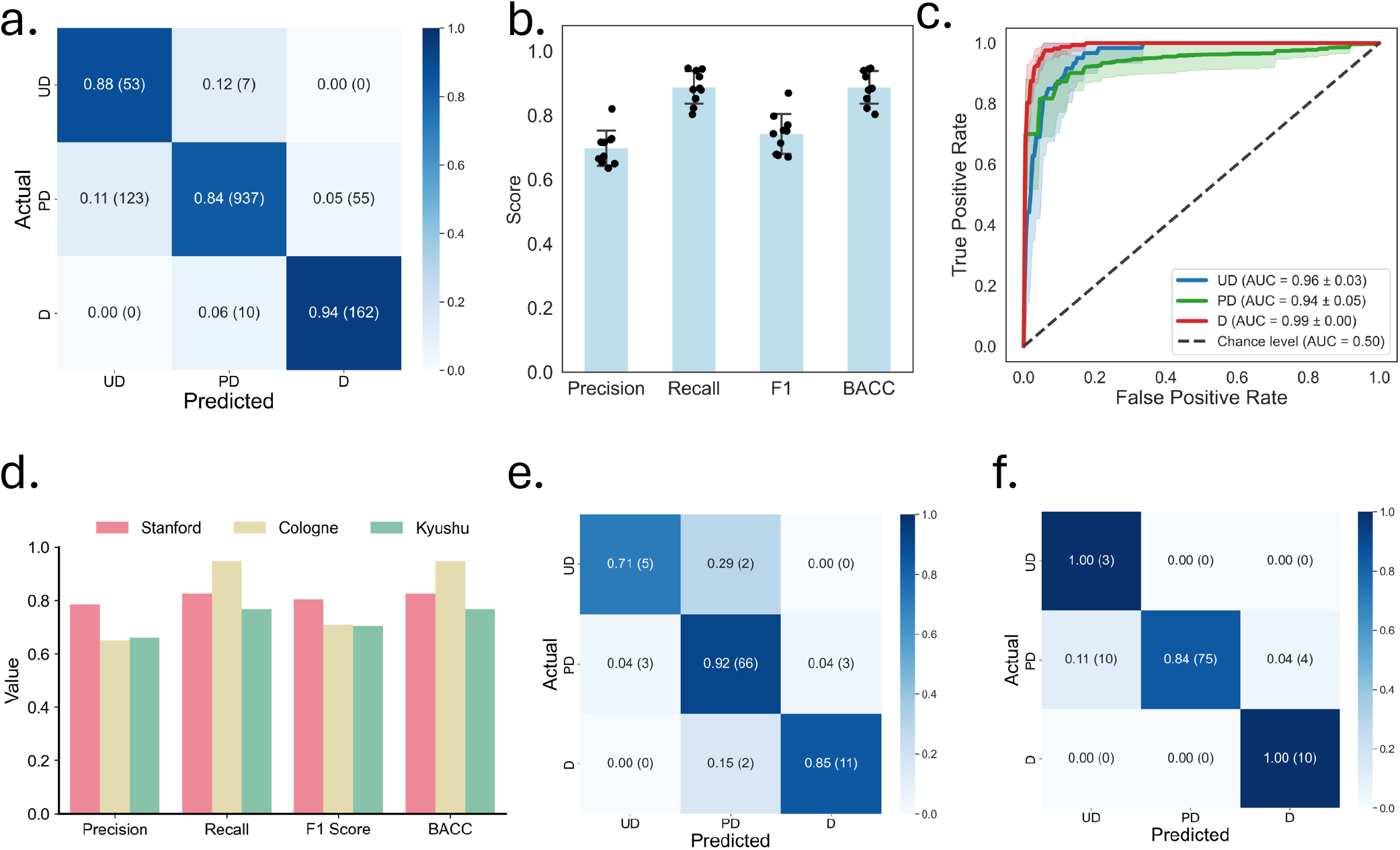
Classification performance for grade of neuroblastic differentiation in tumors of the NB and GNB-n categories. **(a-c)** Performance from ten-fold cross-validation in the COG cohort: (a) confusion matrix showing predictions pooled across the validation sets from all ten folds.; (b) macro-averaged precision, recall, F1 score, and balanced accuracy, with each dot representing the performance of an individual fold. Bars indicate the mean values across the folds, and error bars represent the standard deviation; and (c) per-class receiver operating characteristic curves, with shaded areas representing the standard deviation across folds. **(d)** Classification performance in the independent confirmatory cohorts. **(e-f)** Confusion matrices for the (e) Stanford and (f) Cologne cohorts. D, differentiating; U-P, undifferentiated or poorly differentiated; BACC, balanced accuracy; AUC, area under the receiver operating characteristic curve.

**Table 3.** Distribution of tumors in the NB and GNB-n categories by grade of neuroblastic differentiation and mitosis-karyorrhexis index across cohorts.

| Cohort | Grade subtype |  |  |  | MKI |  |  |
|---|---|---|---|---|---|---|---|
|  | UD | PD | D | UE | Low-or-intermediate | High | UE |
| COG-pNT | 60 | 1,115 | 172 | 0 | 1,148 | 198 | 1 |
| Stanford-NB | 7 | 72 | 13 | 0 | 80 | 12 | 0 |
| Cologne-pNT | 3 | 89 | 10 | 6 | 92 | 7 | 9 |
| Kyushu-pNT | 3 | 109 | 2 | 0 | 106 | 6 | 2 |
| **Total** | **73** | **1,385** | **197** | **6** | **1,426** | **223** | **12** |

UD: undifferentiated; PD: poorly differentiated; D: differentiating; UE: unevaluable; MKI, mitosis-karyorrhexis index. Low and intermediate MKI categories were combined into a single category for model development and evaluation.

To evaluate generalizability, we validated the model on NB and GNB-n cases from the three independent confirmatory cohorts: Stanford, Cologne, and Kyushu (**Table 3**). In the Stanford cohort, the model achieved a precision of 0.78, recall of 0.83, F1 score of 0.80, and balanced accuracy of 0.83 (**Figs. 3d-e**). The AUROC values for undifferentiated, poorly differentiated, and differentiating tumors were 0.96, 0.93, and 0.98, respectively (**Supplementary Fig. 2a**). Similarly, in the Cologne cohort, the model achieved a precision of 0.65, recall of 0.95, F1 score of 0.71, and balanced accuracy of 0.95 (**Figs. 3d, f**). The corresponding AUROC values were 0.99, 0.91, and 0.99 for undifferentiated, poorly differentiated, and differentiating tumors, respectively (**Supplementary Fig. 2b**). In the Kyushu cohort, the model achieved a precision of 0.66, recall of 0.77, F1 score of 0.70, and balanced accuracy of 0.77 (**Fig. 3d and Supplementary Fig. 2c**). Notably, the Kyushu cohort included only two tumors classified as differentiating, both of which were correctly predicted by our model (**Supplementary Fig. 2c**). The relatively lower performance observed in the Kyushu cohort was primarily attributable to misclassification of two of the three undifferentiated tumors as poorly differentiated. The corresponding AUROC values in the Kyushu cohort were 0.99, 0.97, and 1.00 for undifferentiated, poorly differentiated, and differentiating tumors, respectively (**Supplementary Fig. 2d**).

### Predictive Performance for MKI class in tumors of the NB and GNB-n Categories

While most existing approaches for H&E image analysis focus primarily on identifying mitotic figures, they often neglect karyorrhectic cells, despite their critical role in neuroblastoma diagnosis. To address this gap, we next developed a dedicated deep-learning model to predict MKI class from histology images of tumors in the NB and GNB-n categories.

According to the INPC system, tumors are classified into three MKI classes: Low (<100/5,000 tumor cells), Intermediate (100-200/5,000 tumor cells), and High (≥200/5,000 tumor cells)^1–4^. In the NB-PathAssist, however, we distinguished only two MKI classes, Low-or-intermediate vs. High MKI, due to the limited prognostic contribution of Intermediate MKI alone, whereas High MKI has strong adverse prognostic implication by itself^15,17^ (**Table 3**). To identify mitotic and/or karyorrhectic (MK) tumor cells at a single-cell resolution, we implemented a pathologist-in-the-loop framework (**Fig. 4a**). Specifically, we first applied a pre-trained deep learning model to segment nuclei from H&E images^18^, after which each cell was manually annotated into one of three categories: (1) mitotic and/or karyorrhectic (MK) tumor cells, (2) non-MK tumor cells, and (3) non-tumor cells.

**Figure 4.**
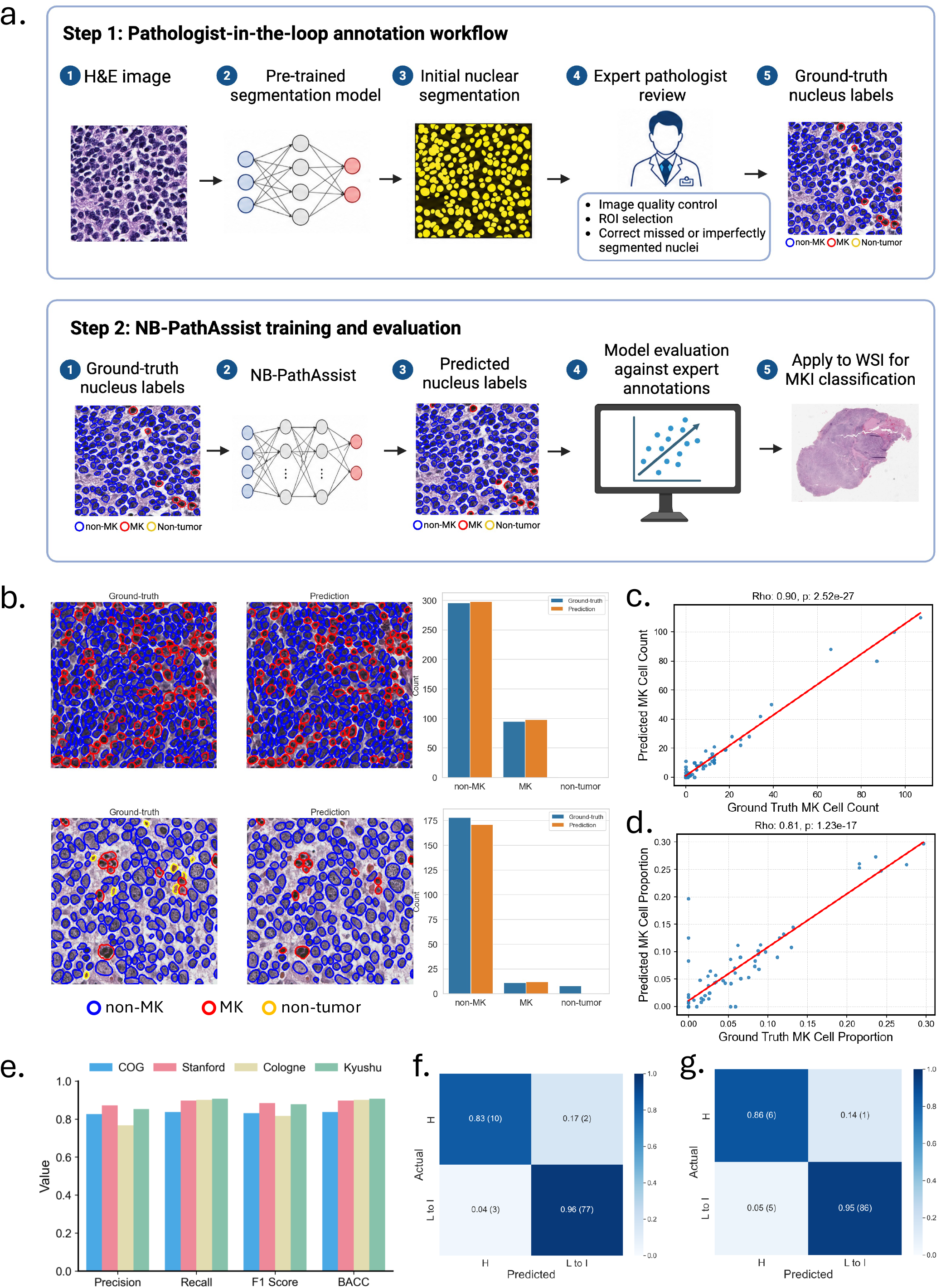
Classification performance for MKI class prediction. **(a)** Pathologist-in-the-loop workflow for training a deep-learning model to detect mitotic and karyorrhectic (MK) cells from H&E-stained histology images. Image was created using BioRender. **(b)** Ground-truth and predicted cell classes in two representative tumor regions, with corresponding cell counts. **(c-d)** Correlation between ground-truth and predicted (c) absolute counts and (d) proportions of MK cells per image. **(e)** MKI classification performance on whole-slide histology images across different cohorts. **(f-g)** Confusion matrices for MKI class prediction on whole-slide images from the (f) Stanford and (g) Cologne cohorts. MKI, mitosis-karyorrhexis index; MK, mitotic-karyorrhectic; H, high MKI; L-I, low or intermediate MKI.

To train the model, we annotated 10,415 cells from 72 images of the neuroblastoma cases within regions selected by expert pathologists from the study cohort, including 7,077 non-MK tumor cells, 793 MK tumor cells, and 2,545 non-tumor cells (**Fig. 4b**). We performed 5-fold cross-validation using 80% of the images for training and hyperparameter tuning and the remaining 20% for held-out evaluation. For independent testing, we additionally annotated 22 images from the Stanford cohort, including 3,353 non-MK cells, 212 MK cells, and 69 non-tumor cells.

In the validation set of the study cohort, the predicted MK cell counts per image were significantly correlated with ground truth (**Figs. 4b-c**, Spearman r = 0.90, P = 2.52 × 10^-27^), and the proportion of MK cells among tumor cells also showed significant correlation (**Fig. 4d**, Spearman r = 0.81, P = 1.23 × 10^-17^). Consistent results were observed in the independent Stanford cohort, where predicted MK cell counts correlated well with ground truth (Spearman r = 0.85, P = 4.14 × 10^-7^), as did the proportion of MK cells (Spearman r = 0.78, P = 1.89 × 10^-5^) (**Supplementary Figs. 3a-c**).

We then applied the best-performing model from five-fold cross-validation to whole-slide histology images of all evaluable NB and GNB-n tumor sections from the study cohort (i.e., COG) and the independent confirmatory cohorts (i.e., Stanford, Cologne, and Kyushu), comprising a total of 1,649 tissue sections. In the COG cohort, the model achieved a macro-averaged precision of 0.83, recall of 0.84, F1 score of 0.83, and balanced accuracy of 0.84 (**Fig. 4e**). In the Stanford cohort, the model achieved a macro-averaged precision of 0.87, recall of 0.90, F1 score of 0.88, and balanced accuracy of 0.90 (**Figs. 4e-f**). The AUROC score of identifying high MKI tumors in the Stanford cohort was 0.99 (**Supplementary Fig. 3f**). A similar performance was observed in the Cologne cohort, with a precision of 0.77, recall of 0.90, F1 score of 0.81, balanced accuracy of 0.90, and AUROC of 0.99 (**Figs. 4e, g and Supplementary Fig. 3g**). In the Kyushu cohort, the model achieved a precision of 0.85, recall of 0.91, F1 score of 0.88, balanced accuracy of 0.91, and AUROC of 0.91 (**Fig. 4e and Supplementary Figs. 3h-i**). Overall, these results indicate that our model can faithfully predict MKI class of NB and GNB-n tumors.

### Performance for Prognostic Stratification

According to the INPC system, pNTs are classified into Favorable Histology (FH) or Unfavorable Histology (UH) groups (**Table 1)**. Tumors in the NB and GNB-n categories are classified into FH or UH based on the age-linked evaluation of grade of neuroblastic differentiation and MKI. In contrast, GNB-i and GN are always classified as FH. Patients with FH tumors have significantly better survival outcomes than those with UH tumors^3,17,19^.

To demonstrate the clinical utility of our AI framework, we evaluated its ability to classify tumors as FH or UH by integrating patient age and the predictions of tumor category, grade of neuroblastic differentiation, and MKI from whole-slide histology images. For GN or GNB-i cases misclassified as NB or GNB-n (see **Fig. 2**), we additionally applied the grade and MKI models to assign the corresponding prognostic group. Across the validation datasets of the COG cohort, the resulting prognostic classification achieved a macro-averaged precision of 0.93, recall of 0.94, F1 score of 0.93, and balanced accuracy of 0.94 (**Figs. 5a-b**).

**Figure 5.**
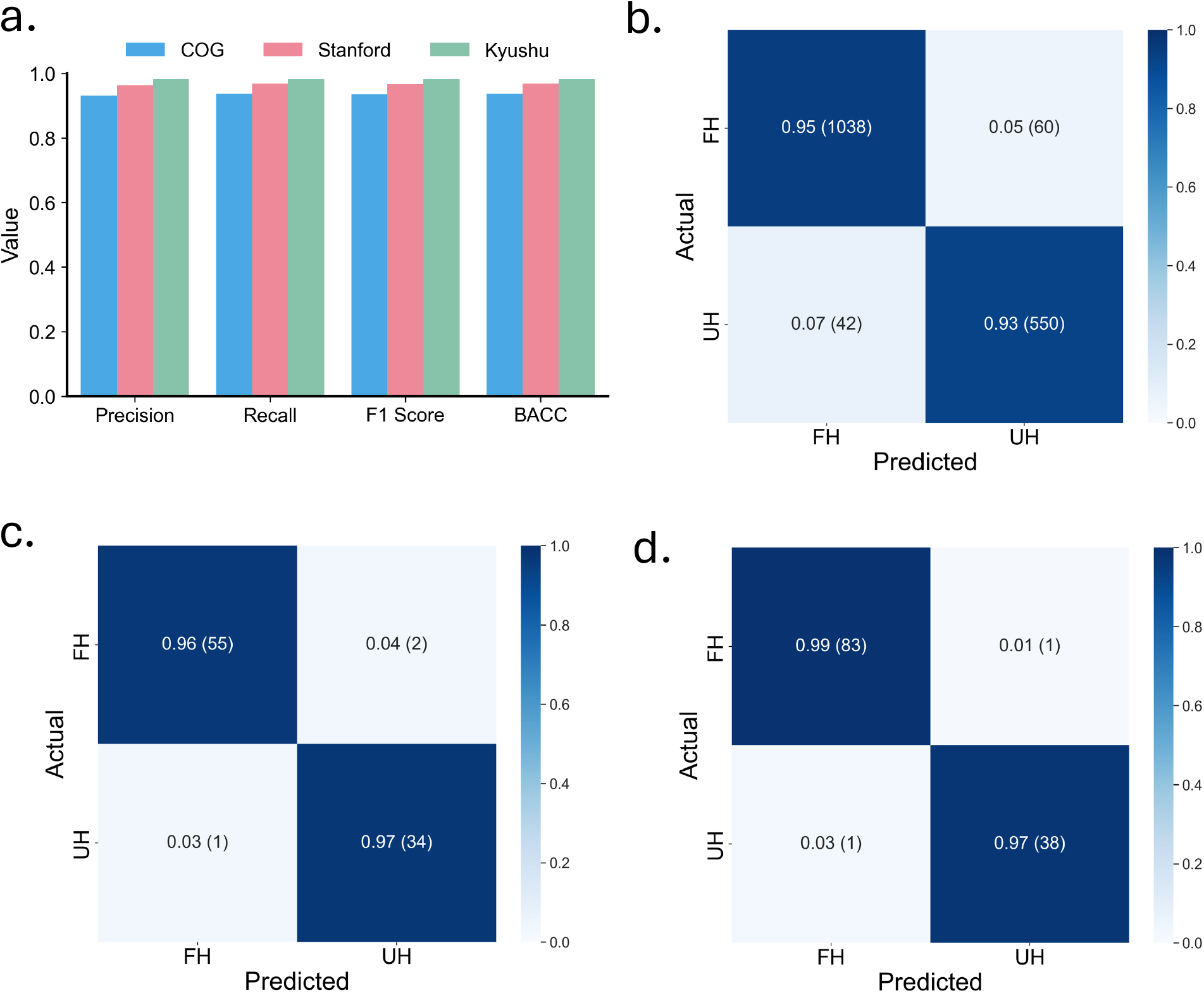
Model performance for prognostic stratification. **(a)** Classification performance for prognostic stratification across cohorts**. (b-d)** Confusion matrices for classifying favorable versus unfavorable histology in the (b) COG, (c) Stanford, and (d) Kyushu cohorts. FH, favorable histology; UH, unfavorable histology; BACC, balanced accuracy.

We then validated the model performance in the Stanford and Kyushu cohorts, for whom patient age information was available. In the Stanford cohort, the model achieved a macro-averaged precision of 0.96, recall of 0.97, F1 score of 0.97, and balanced accuracy of 0.97 (**Figs. 5a, c**). In the Kyushu cohort, the model achieved a precision of 0.98, recall of 0.98, F1 score of 0.98, and balanced accuracy of 0.98 (**Figs. 5a, d**). These findings underscore the utility of our models in prognostic stratification from routinely collected H&E images.

### An online digital platform for automated neuroblastic tumor analysis

To enable the community to use our models, we developed an online computational platform, NB-PathAssist (https://nb-ai.stanford.edu), for end-to-end analysis of pNT histology images (**Figs. 6a-b**). Through the platform, users can upload images of tumor sections stained with H&E directly from their local systems, after which the software automatically performs data preprocessing, feature extraction, model inference, and prediction of tumor category and further histologic features. The platform then generates a comprehensive textual report summarizing the prognostic classification (FH vs. UH) of the tumor.

**Figure 6.**
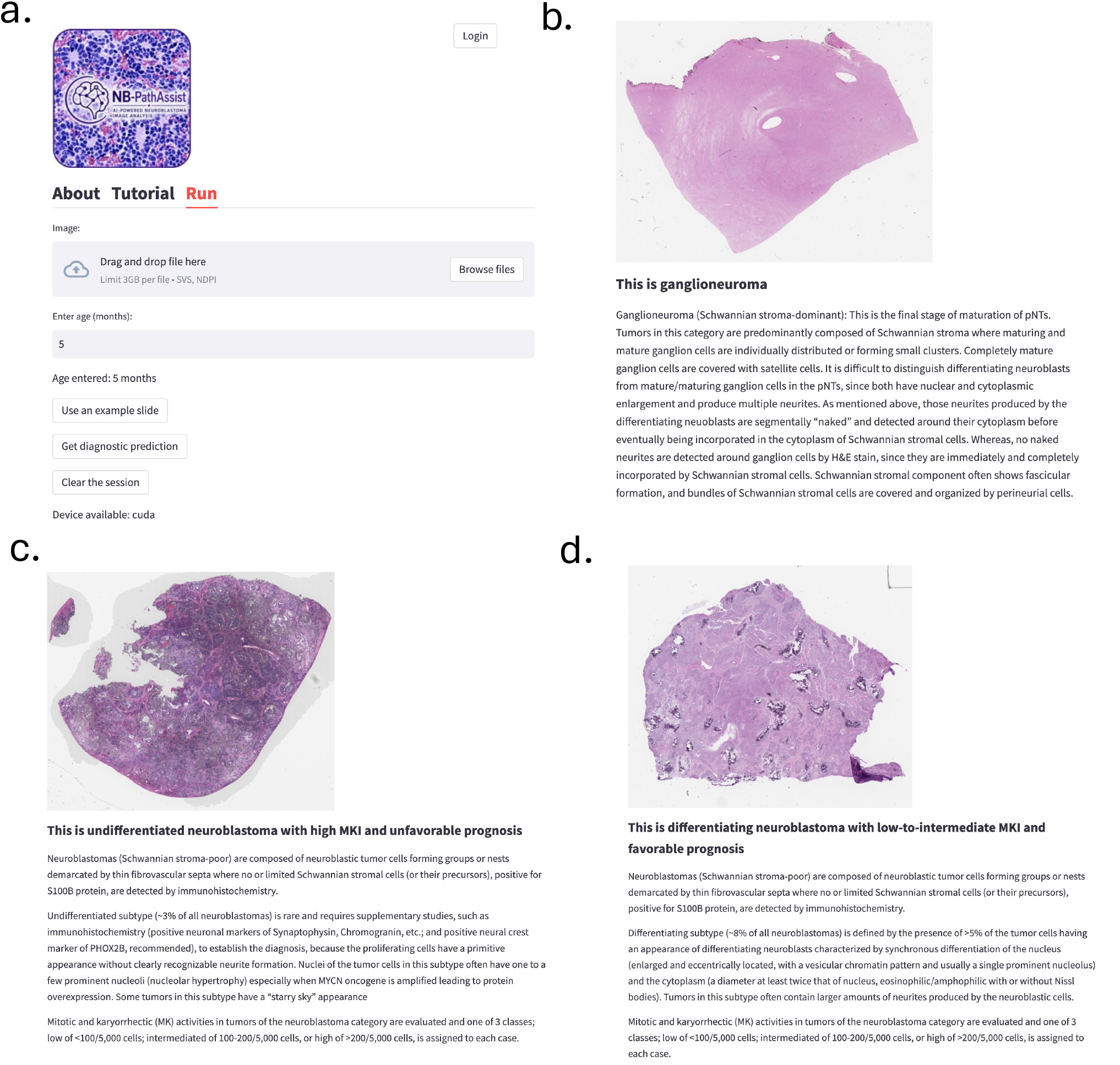
NB-PathAssist: an online computational tool for automated neuroblastic tumor classification. (**a**) Screenshot of user interface. (**b-d**) Example predictions for (b) a ganglioneuroma; (c) a neuroblastoma with high MKI and undifferentiated subtype with unfavorable prognosis, and (d) a neuroblastoma with low-to-intermediate MKI and differentiating subtype with favorable prognosis.

## Discussion

Peripheral neuroblastic tumors (pNTs), including neuroblastoma, ganglioneuroblastoma, and ganglioneuroma, make the most common extra-CNS solid tumors in pediatric age group^1–4^. It is also well known that pNTs presents a wide range of behaviors, such as spontaneous regression, tumor differentiation/maturation, and aggressive progression refractory to intensive treatment^20,21^. Historically, clinical outcomes of the patients with this disease are predicted by so-called prognostic factors. Histopathology according to the INPC, one of those prognostic factors, has been considered highly significant for predicting prognosis of the patients by distinguishing the FH group (5-year Event-free Survival of 97.3+/-1.3% and 5-year Overall survival of 96.0+/-0.7%) and the UH group (EFS of 51.1+/-1.9%; OS of 61.5+/-1.8%) with EFS Hazard Ratio (95% confidence interval) of 4.41 (3.85, 5.03) and OS Hazard Ratio of 10.50 (8.44, 13.04)^3,7^.

Traditionally, prognostic distinction among pNTs is determined by pathologists through review of histopathologic images. However, this process demands substantial expertise from pathologists and is subject to intra- and inter-observer variability^22^, a challenge exacerbated by the global shortage of specialized pathologists. In the past decade, deep-learning models trained on whole-slide histology images of tumor tissues have shown promise in supporting cancer diagnosis and prognosis^9,10^. However, existing models are largely trained on data from adult disease, with limited applications on rare pediatric disease. In 2009, Kong et al. developed a hierarchical classification algorithm that utilized textural patterns extracted from multi-resolution image segmentation to predict the grade of neuroblastic differentiation. Their classifier was trained on 387 image tiles derived from three whole-slide images^23^. Subsequently, Liu et al. performed nucleus segmentation and extracted morphological and intensity features associated with favorable or unfavorable prognosis using 563 WSIs from 107 patients^24^. Although these studies have demonstrated the potential of computational pathology for neuroblastoma diagnosis and prognostic assessment, most have been evaluated in relatively small cohorts, in part because of the limited availability and accessibility of digitized histopathology data. This challenge is further complicated by the imbalanced distribution of key histopathologic features: fewer than 10% of tumors are classified as differentiating, and only approximately 20% exhibit high MKI^15,16^. Models developed on such datasets may therefore be susceptible to overfitting and may not generalize across independent patient populations.

In this study, we assembled the largest multi-institutional whole-slide image cohort of peripheral neuroblastic tumors to date, comprising 2,030 tissue sections collected from institutions across multiple geographic regions. To improve the representation of less common histopathologic subgroups, we enriched the cohort for differentiating neuroblastoma and tumors with high MKI. Model performance was rigorously evaluated through internal cross-validation and independent validation using geographically diverse confirmatory cohorts spanning multiple continents. To improve prediction robustness and reduce variability associated with individual models, we employed an ensemble approach in which models trained across ten cross-validation folds were independently applied to each case in the confirmatory cohorts, and their predictions were averaged to generate the final classification.

A key methodological advancement of our approach is the quantification of MKI through single-cell segmentation and classification. Specifically, we developed a deep-learning model for simultaneous nucleus segmentation and cell-type classification in H&E-stained images using annotations generated by expert pathologists. We manually annotated more than 14,000 nuclei and classified each cell into one of the three categories: mitotic/karyorrhectic (MK) tumor cells, non-MK tumor cells, or non-tumor cells. The trained model was then applied to whole-slide images to quantify the proportion of MK cells among tumor cells, and MKI class was assigned according to established pathological standards. Compared with existing approaches that directly predict MKI from patch-level features using histology foundation models and multiple-instance learning^14^, our cell-based strategy offers several advantages. First, our single-cell approach closely mirrors the pathological definition of MKI, which is based on the frequency of mitotic and karyorrhectic cells rather than on global tissue morphology. By explicitly identifying and quantifying the cellular events that determine MKI, the model provides a biologically and clinically interpretable measurement that can be directly compared with established diagnostic criteria. Second, cell-level supervision provides substantially denser training signals than slide-level labels. This is particularly advantageous for detecting relatively rare MK cells and may reduce reliance on indirect morphological correlates that could vary across institutions or staining conditions. In contrast, a slide-level multi-instance learning model may achieve accurate classification by exploiting correlated but non-causal histological features, making its predictions more difficult to interpret and potentially less robust to distribution shifts. Finally, the cell-based approach produces intermediate outputs, including segmented nuclei, cell classifications, and MK-cell proportions, that can be visually inspected and independently audited by pathologists, thereby increasing the transparency of the resulting MKI assessment.

One limitation of this study is that the model did not distinguish tumors with intermediate MKI from those with low MKI. Instead, these two categories were combined into a single low/intermediate MKI group. This design option was motivated by previous observations that intermediate MKI alone has limited prognostic significance^15,17^. However, under the INPC system (**Table 1**), MKI can affect histologic prognostic classification in specific clinical contexts. In patients aged >1.5 to 5 years with differentiating tumors, those with intermediate MKI are classified as having Unfavorable Histology, whereas those with low MKI are classified as Favorable Histology. In our study cohort, the inability to distinguish low from intermediate MKI affected only 5 out of 1,690 (0.3%) tumors: these cases were classified as Unfavorable Histology based on the original COG diagnosis but as Favorable by NB-PathAssist. Despite this limitation, NB-PathAssist reached excellent performance in classifying the INPC prognostic group across geographically diverse patient cohorts, with an average precision of 0.96 (std: 0.03), recall of 0.96 (std: 0.02), F1 score of 0.96 (std: 0.03) and balanced accuracy of 0.96 (std: 0.02).

In summary, our study combines methodological innovation with the largest-to-date neuroblastic tumor cohorts to advance diagnostic and prognostic strategies for these rare pediatric tumors. We demonstrate that neuroblastoma diagnostic categories can be accurately predicted from H&E-stained histology images using deep learning. Although this study focuses on neuroblastic tumors, the proposed deep-learning workflow and model architectures are broadly adaptable and may support AI-assisted histopathologic diagnosis across other pediatric and adult cancers.

## Methods

### Cohort characteristics and ethics

The study was approved by the Stanford’s Institutional Review Board (IRB) (#46426, #73907).

COG-pNT: The COG collection of peripheral neuroblastic tumors (pNTs) contains 1,692 tumor sections from 1,692 patients across the United States, Canada, Australia, and New Zealand. Of the cohort, 1,264 patients were diagnosed with neuroblastoma (NB, Schwannian stroma-poor), 242 patients were diagnosed with ganglioneuroblastoma, intermixed (GNB-i, Schwannian stroma-rich), and 103 with ganglioneuroma (GN, Schwannian stroma-dominant). In addition, 83 patients were diagnosed with ganglioneuroblastoma, nodular (GNB-n, composite, Schwannian stroma-rich/stroma-dominant and stroma-poor). The median age at diagnosis was 22 months, with an interquartile range of 9 to 50 months.

Stanford-NB: The Stanford cohort contains 92 tissue sections of neuroblastoma from 81 patients treated at Stanford Medicine Children’s Health, USA. The median age at diagnosis was 14 months, with an interquartile range of 5 to 27 months.

Cologne-pNT: The Cologne cohort contains 121 pNT tissue sections from 120 patients treated at the University Children’s Hospital of Cologne, Germany. Of the cohort, 102 tissue sections were NB, 7 were GNB-i, 6 were GN, and 6 were GNB-n. Of the NB and GNB-n cases, tumor grade was evaluable in 102 tissue sections, whereas MKI was evaluable in 98 sections. Age information was not available for this cohort.

Kyushu-pNT: The Kyushu cohort contains 125 pNT tissue sections from 125 patients treated at Kyushu University Hospital, Japan. The cohort comprises NB cases (n = 109), GNB-i (n = 7), GN (n = 4), and GNB-n (n = 5). Of the NB and GNB-n cases, tumor grade was evaluable in all 114 tissue sections, whereas MKI was evaluable in 112 sections. The median age at diagnosis is 11 months, with an interquartile range of 7 to 24 months.

### Whole-slide image preprocessing

Whole-slide histology images (WSIs) from the COG-pNT, Stanford-NB and Kyushu-pNT cohorts were acquired in SVS format using an Aperio slide scanner, whereas WSIs from the Cologne-pNT cohort were acquired in NDPI format using a Hamamatsu NanoZoomer scanner. All images were subsequently downsampled to 20× magnification for downstream analysis. Background regions were removed using the Otsu thresholding algorithm^25^. A board-certified pathologist (H.S.) reviewed the images, and those with insufficient evaluable tumor tissue, complete necrosis, or poor focus were excluded from downstream analysis. Given that each WSI typically contains millions of pixels, we divided each image into non-overlapping tiles of size 256 x 256 pixels (equivalent to 128 μm x 128 μm). Tiles containing over 20% background area were excluded from further analysis. Each remaining tile was transformed into a linear feature vector of size 1 1,024 using UNI, a recently developed foundation model for histology images^11^. For model training, we randomly selected *N* tiles from each WSI, with *N* set to 4,000 based on our prior benchmarks^10^.

### Model architecture, training and evaluation for predicting tumor category and grade of differentiation

According to the INPC/WHO framework, neuroblastic tumor category is defined by the degree of neuroblastic differentiation and Schwannian stromal development, whereas neuroblastoma grade is further determined by the extent of tumor cell differentiation^1–3^. These diagnostic features can be spatially heterogeneous across a whole-slide image, and in nodular or composite tumors, the highest-grade component may drive the final classification. Therefore, slide-level prediction requires integration of heterogeneous histologic patterns across multiple tumor regions rather than classification based on isolated image patches.

To address the spatial heterogeneity of neuroblastic tumors, we developed an archetype-guided SummaryMixing architecture, termed Archetype-SM, for WSI-level prediction of tumor category and grade of differentiation. Each WSI was first divided into image patches, and patch-level morphologic features were extracted using a feature extraction module. To reduce redundancy while preserving representative histologic diversity, patch features were compressed into *T* 100 archetype features, denoted as *X* = {x*_1_*,…, x*_T_*}, where each archetype captures a representative morphologic pattern within the slide. These archetype features were then used as input tokens for Archetype-SM.

Instead of applying conventional self-attention, which explicitly computes pairwise interactions among all input tokens and scales quadratically with token number, Archetype-SM uses a SummaryMixing-based token-mixing module^26^. SummaryMixing provides a linear-complexity alternative to self-attention by first computing a global summary vector across all tokens and then combining this global context with each token-specific representation. In our implementation, the slide-level summary vector s was computed across all archetype tokens and injected into each archetype-specific transformation, enabling efficient modeling of global histologic composition while maintaining linear computational complexity.

Formally, the operation is defined as:

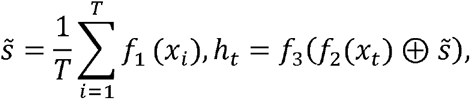

Where ⊕ represents concatenation, and *f*_1_ f_2_ and *f_3_* are dense layers followed by GeLU activation. By combining archetype compression with linearized transformer aggregation, Archetype-SM provides a scalable approach for learning slide-level representations from morphologically heterogeneous tumors. This is particularly relevant for neuroblastic tumor classification, where diagnostic category and grade depend on the relative representation of neuroblastic differentiation, Schwannian stromal development, and high-grade tumor components across the slide.

For model training and internal cross-validation, we split data from the study cohort into 10 folds. Each fold included 90% of the data for global training and 10% for testing. Within each global training set, we further split the data into 90% for training and 10% for hyperparameter tuning. This tuning set was used to determine the optimal stopping point (i.e., the number of training epochs) for model saving. The model with optimal performance on the tuning set was then applied to the testing cohort for evaluation. We used cross-entropy as the objective function and optimized the model with the Adam optimizer^27^.

To evaluate the model’s predictive performance on independent cohorts, we utilized an ensemble strategy, where models trained on each fold of the study cohort were separately applied to each image, and the class probabilities predicted by these models were averaged to obtain the final prediction. The performance metrics included macro-F1 score, precision, recall, and the area under the receiver operating characteristic curve (AUROC).

### Model architecture, training and evaluation for predicting MKI class

In routine pathological practice, the mitosis-karyorrhexis index (MKI) is determined by the proportion of mitotic and karyorrhectic nuclei relative to the total number of neuroblastic nuclei. To align our AI framework with established pathological assessment methodology, we developed a deep-learning pipeline for nuclei segmentation and classification in neuroblastic tumor histology images.

We first utilized a recently established deep-learning model to segment cell nuclei from H&E-stained images^18^. The segmentation results were independently reviewed by H.S. and Y.S., and incorrectly segmented or missed nuclei were manually corrected and annotated using QuPath (version 0.6.0).

Each nucleus was subsequently assigned to one of three categories: (1) mitotic and/or karyorrhectic (MK) cell, (2) non-MK neuroblastic cell, or (3) non-tumor cell.

A transformer-based model was then trained using nucleus-level image embeddings to classify nuclei into these categories. Model training was performed with a learning rate of 5 × 10⁻ □, a batch size of 16, and a maximum of 400 epochs.

## Acknowledgement

We are grateful for the generous support from the U.S. Food and Drug Administration (FDA) (Contract No. 75F40123C00103) and Children’s Oncology Group (NCTN Operations Center Grant U10CA180886, COG Biospecimen Bank Grant U24CA196173, NCTN Statistics & Data Center Grant U10CA180899). Y.Z. was further supported by the NIH/NCI Pathway to Independence award (1K99CA293249). This work was further supported by the Deutsche Forschungsgemeinschaft (DFG) as part of the CRC1588 (grant 493872418 to M.F.) and CRC1399 (grant 413326622 to L.W. and M.F.), the Bruno und Helene Joester Stiftung, the Ministry of the Interior of Northrine-Westphalia, the Ministry of Culture and Science of the State of Northrhine-Westphalia through CANTAR (to M.F.), and by Förderverein für krebskranke Kinder e.V. Köln (to M.F.). The content is solely the responsibility of the authors and does not necessarily represent the official views of the funding agencies.

## Figure Legends

**Supplementary Figure 1.** Classification performance for tumor category by treating GNB-i and GN as separate class. **(a-c)** Performance in the COG cohort: (a) confusion matrix; (b) macro-averaged precision, recall, F1 score, and balanced accuracy from ten-fold cross-validation; and (c) per-class receiver operating characteristic curve, with shaded areas representing the standard deviation across cross-validation folds. **(d-f)** Classification performance in the confirmatory cohorts: (d) confusion matrix; € macro-averaged precision, recall, F1 score, and balanced accuracy; and (f) per-class receiver operating characteristic curves. NB, neuroblastoma; GNB-i, ganglioneuroblastoma, intermixed; GN, ganglioneuroma; GNB-n, ganglioneuroblastoma, nodular; BACC, balanced accuracy; AUC, area under the receiver operating characteristic curve.

**Supplementary Figure 2.** Classification performance for grade of neuroblastic differentiation in independent confirmatory cohorts. **(a-b)** Per-class receiver operating characteristic curves in the (a) Stanford and (b) Cologne cohorts. **(c)** Confusion matrix for grade prediction in the Kyushu cohort. **(d)** Per-class classification performance in the Kyushu cohort. D, differentiating; U-P, undifferentiated or poorly differentiated; AUC, area under the receiver operating characteristic curve.

**Supplementary Figure 3.** Classification performance for MKI class in independent confirmatory cohorts. **(a)** Ground-truth and predicted cell classes in two representative tumor regions from selected images of the Stanford cohort, with corresponding cell counts. **(b-c)** Correlation between ground-truth and predicted (b) absolute counts and (c) proportions of MK cells per image. **(f-g)** Receiver operating characteristic (ROC) curves showing the performance of the model in identifying tumors with high MKI in the (f) Stanford and (g) Cologne cohorts. **(h)** Confusion matrix for MKI class prediction in the Kyushu cohort. **(i)** ROC curve showing the performance of the model in identifying tumors with high MKI in the Kyushu cohort. MKI, mitosis-karyorrhexis index; MK, mitotic and karyorrhectic; AUC, area under the receiver operating characteristic curve.

