## Supplementary figures and images for "Digital pathological classification of peripheral neuroblastic tumors across multi-continent cohorts"

### Supplementary Figures

#### Supplementary Figure 1

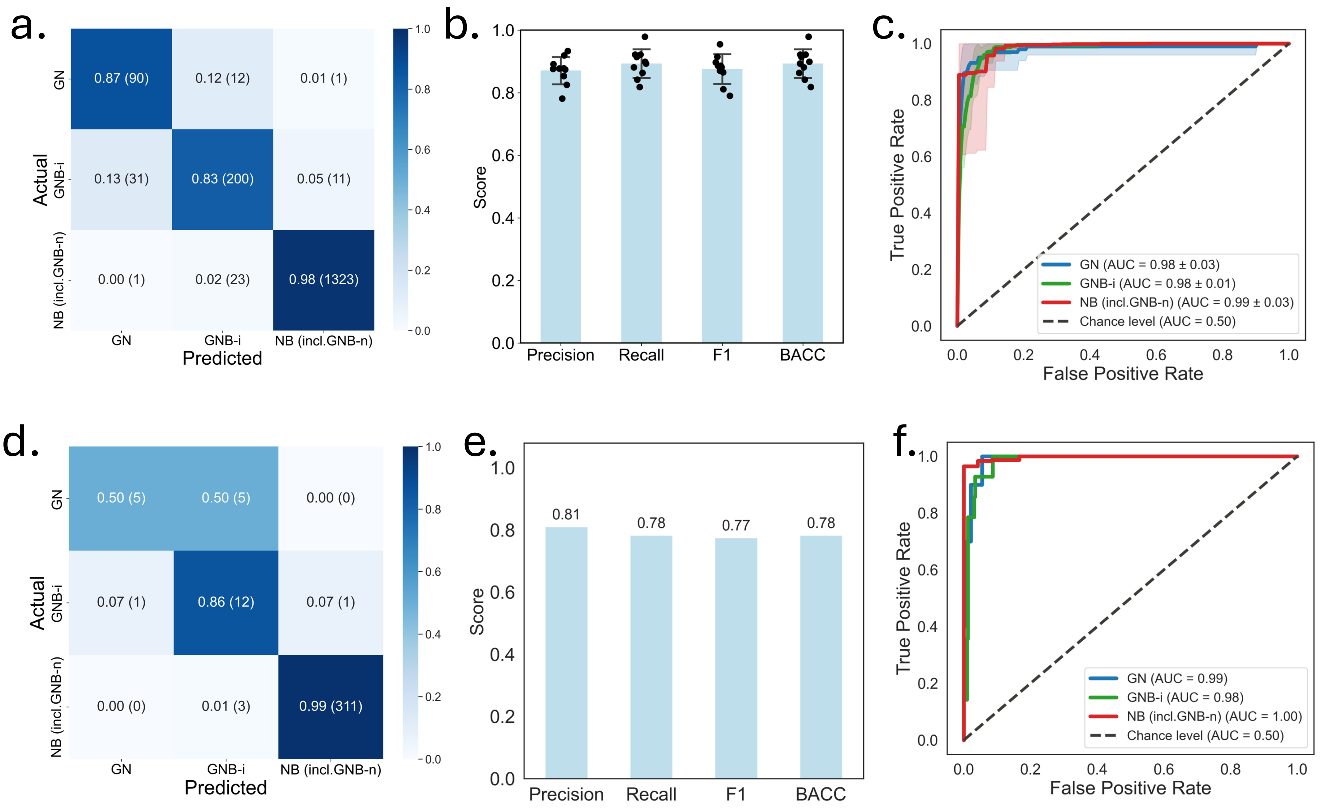

**Supplementary Figure 2**

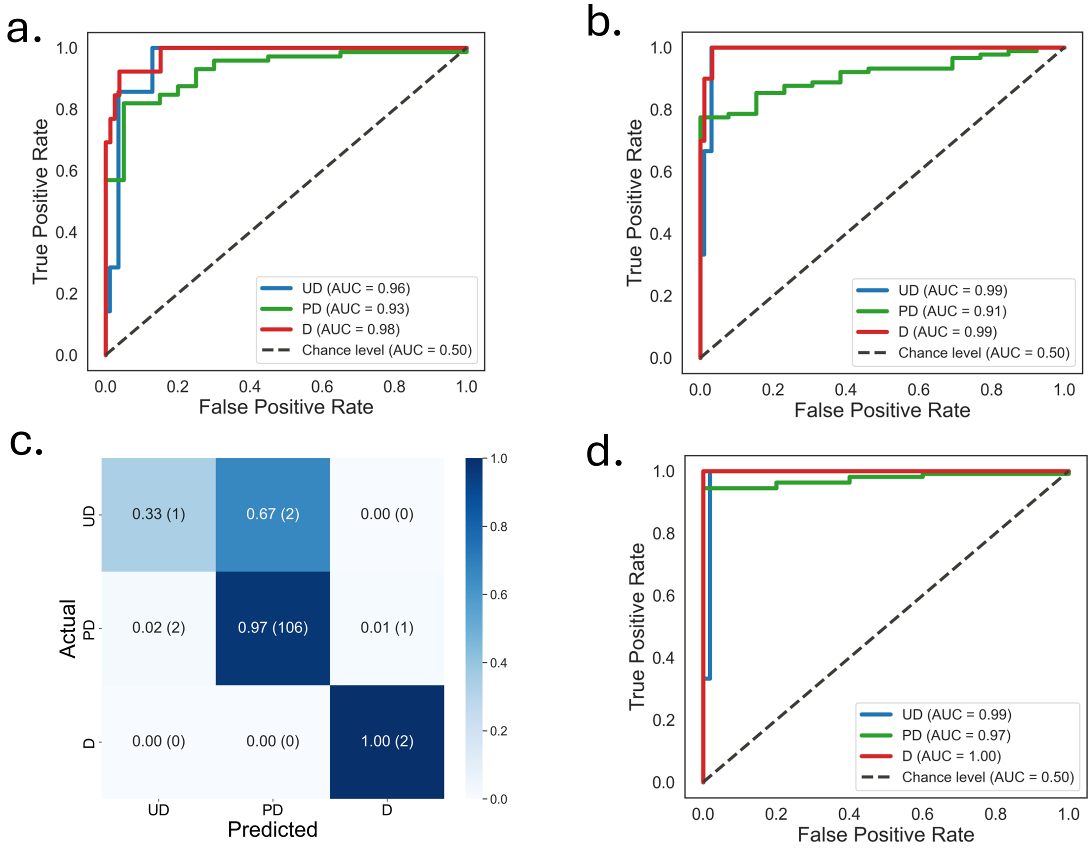

#### Supplementary Figure 3

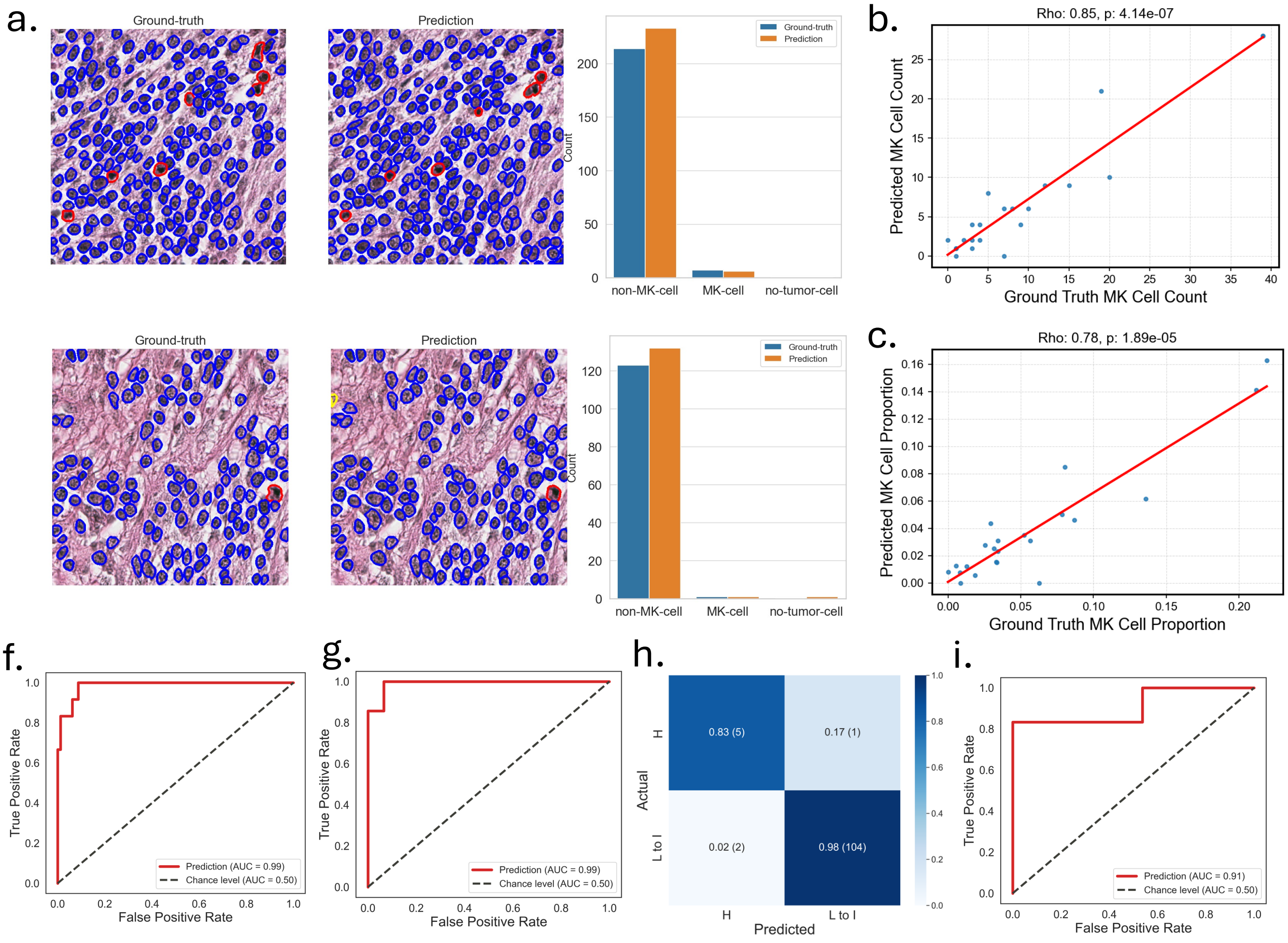
